# Railway Catenary Sparking as a Source of Toxic Copper Ultrafine Particles: Evidence from Realistic In Vitro Inhalation Exposure

**DOI:** 10.64898/2026.05.07.723476

**Authors:** J. Becker, J. Pantzke, S. Offer, A. Das, A. Mudan, C. Neukirchen, T. Streibel, T. Adam, M. Sklorz, S. Di Bucchianico, R. Zimmermann

## Abstract

Railway catenary sparking generates airborne ultrafine particles (UFPs) that may pose health risks due to their metallic composition and ability to penetrate deep into the alveolar region of the lungs. Copper, widely used in wires and pantographs, is a major component of these emissions, making copper-rich particles common in railway environments such as subways. However, exposure levels and health impacts remain poorly characterized, and localized hotspots may represent an underrecognized risk in densely populated areas. This study investigated the toxicity of copper UFPs under realistic dosimetry and deposition conditions. Copper UFPs were generated using a spark discharge generator and applied to two *in vitro* lung models: a 3D co-culture of Calu-3 epithelial cells, THP-1-derived macrophages, and EA.hy926 endothelial cells, and a monoculture of A549 alveolar epithelial cells. Cells were exposed at the air–liquid interface (ALI) using an automated platform to mimic inhalation exposure and UFPs deposition. Copper deposition ranged from 6.5 to 41 ng/cm^2^, within occupationally relevant levels. A549 cells showed cytotoxic responses consistent with previous studies, whereas the 3D co-culture model revealed broader adverse effects, including inflammation, impaired epithelial barrier integrity, oxidative stress, and early DNA damage. Inflammatory activation also differed between models: A549 cells mainly exhibited transcriptional responses, while the 3D model showed significant secretion of IL-6 and IL-8, associated with interferon signaling. These findings highlight the potential health risks of copper UFPs from railway systems and emphasize the need for improved characterization of UFP exposure in environmental and occupational railway settings.

## Introduction

Unlike tailpipe emissions, non-exhaust emissions arise from mechanical or electrical wear processes such as rail catenary sparking or vehicle braking and therefore exhibit distinct chemical and physical properties. In electrified rail systems, pantograph–catenary interactions can generate substantial quantities of metal-containing particulate matter, with emitted composition strongly influenced by the contact material used (Fruhwirt et al. 2023, Minguillon et al. 2018). As electrification of mainline railways expands and tram and subway networks continue to grow, emissions from these non-exhaust sources are likely to increase. This is of particular concern in densely populated urban areas where exposure potential is high. Accordingly, subway systems have been widely investigated as polluted microenvironments in which metals constitute a major fraction of airborne particulate matter (PM), and *in vitro* studies suggest that metal content is an important determinant of toxicity (Loxham and Nieuwenhuijsen 2019, Vallabani et al. 2023).

Copper and copper alloys are widely used as contact wire materials in European railway systems (Fruhwirt et al. 2023). In addition, PM collected in subway environments consistently contains significant copper fractions, alongside iron-rich wear particles (Bendl et al. 2023, Kim et al. 2010). Long-term exposure to copper-containing particles has been associated with adverse respiratory and cardiovascular outcomes, including chronic obstructive pulmonary disease, pneumonia mortality, and cardiovascular morbidity (Zhang et al. 2021). Experimental studies further indicate that the inhalation toxicity of brake wear particles is partly driven by their copper content (Gasser et al. 2009). For example, copper-enriched brake wear particles induced oxidative stress, inflammation, and disruption of alveolar cellular homeostasis *in vitro* (Figliuzzi et al. 2020, Parkin et al. 2025).

Within the PM size spectrum, ultrafine particles (UFPs; aerodynamic diameter ≤ 100 nm) are of particular concern because they often induce greater biological responses in lung models than larger particles (Karlsson et al. 2009, Midander et al. 2009). UFPs present a distinct inhalation hazard due to their ability to evade upper airway clearance and deposit efficiently in the alveolar regions, where diffusion-driven transport, high surface-area-to-mass ratio, and enhanced surface reactivity promote oxidative stress and reactive oxygen species (ROS) generation (Kwon et al. 2020, Moschini et al. 2023, Schraufnagel 2020). In addition, some UFPs may persist longer in pulmonary tissue and translocate across the air-blood barrier into systemic circulation (Miller et al. 2017). Because air quality standards are largely mass-based (e.g. PM_2.5_, PM_10_), they may insufficiently capture exposures to UFPs, which contribute little to total particulate mass despite high particle number concentrations (Oberdorster et al. 2007).

Substantial toxicological evidence has accumulated on copper and copper oxide UFPs, with more limited epidemiological evidence (Ameh and Sayes 2019). Compared with larger copper particles, nanoscale copper particles show greater redox activity, ion release, and surface reactivity, contributing to distinct toxicological profiles (Midander et al. 2009). Copper-based UFPs are relatively soluble compared with many poorly soluble combustion-derived particles, and intracellular release of dissolved copper ions is considered a major driver of cytotoxic and pro-inflammatory effects observed *in vitro* and *in vivo* (Ahamed et al. 2015, Jing et al. 2015, Moschini et al. 2023). Animal studies further show that pulmonary exposure to copper nanoparticles can induce lung injury and inflammation (Pietrofesa et al. 2021), while chronic exposure may promote immune-cell recruitment and fibrotic remodeling (Lai et al. 2018, Zhang et al. 2024).

At present, *in vitro* toxicity studies of copper UFPs are dominated by submerged exposure designs, in which engineered nanoparticles or copper rich PM samples are suspended in liquid media and applied to cells (Moschini et al. 2023, Parkin et al. 2025). These studies report cytotoxic, inflammatory, and genotoxic effects in human airway models, but exposure doses are often substantially higher than realistic inhalation levels, and particles transformation or agglomeration in suspension remains a major limitation (Midander et al. 2009). Air-liquid interface (ALI) exposure systems provide a more physiologically relevant alternative by enabling direct aerosol deposition onto lung cells while avoiding artifacts associated with submerged dosing (Paur et al. 2011). Even low UFPs doses can induce adverse responses under ALI conditions (Jing et al. 2015). Previous ALI studies with copper nanoparticles reported disturbed metal homeostasis and inflammatory effects in A549 or human bronchial epithelial cells at deposited doses below 1 µg/cm^2^ (Hufnagel et al. 2020, Jing et al. 2015, Kim et al. 2013). However, most studies have relied on monocultures, which do not capture interactions among epithelial, immune, stromal, and endothelial cells that shape pulmonary responses to particle exposure (Hufnagel et al. 2021, Niu and Tang 2022). More advanced co-culture models have shown stronger and more complex inflammatory response than monocultures, including evidence of profibrotic signaling (Pantzke et al. 2022). In addition, inflammatory mediators released in the lung may cross the epithelial barrier and enter circulation, contributing to systemic inflammation, endothelial dysfunction, coagulation changes, and cardiovascular risk (Schraufnagel 2020).

This study addresses key knowledge gaps regarding the toxicity of UFPs generated by rail catenary sparking. A Spark discharge generator (SDG) was used to produce laboratory-generated copper UFPs (SDG-UFP) as a surrogate for catenary-sparking emissions, a well-established method for generating metal nanoparticles for inhalation toxicology studies (Das et al. 2025, Tabrizi et al. 2008). SDG-UFP were assessed in two ALI lung models: (i) an advanced 3D-oriented co-culture comprising bronchial epithelial cells, differentiated macrophages, and vascular endothelial cells (Grytting et al. 2024), and (ii) the widely used A549 alveolar epithelial monoculture model. Endpoints included cytotoxicity, epithelial barrier integrity, cell-type-specific transcriptomic responses, inflammatory and immune-related protein release, oxidative stress markers, and DNA damage. This design enables simultaneous evaluation of effects at the primary deposition site (epithelial and macrophage compartment) and secondary responses in the vascular compartment, representing systemic circulation. The study therefore provides mechanistic insight into how inhaled copper UFP may contribute to both local pulmonary injury and distal systemic effects.

## Material and Methods

### Copper UFP generation and aerosol characterization

Spark discharge generator ultrafine particles (SDG-UFP), were produced and characterized as previously described by (Das et al. 2025). Briefly, particles were generated by a spark discharge generator (PALAS) with copper electrodes in argon atmosphere. Particles were neutralized and passed through a 5 L vessel and a perfluoroalkoxy (PFA) coil to generate the desired particle size and organics were removed by a char coal denuder (Helsatech). After an initial 1:10 dilution, aerosols were directed to the automated ALI cell exposure station (AES, Vitrocell Sytems GmbH, Germany). In a separate sampling line, aerosols were further diluted 1:10 and particle size distribution was measured using a scanning mobility particle sizer (SMPS, TSI).

### TEM imaging

For transmission electron microscopy (TEM) SDG-UFP were deposited onto perforated carbon film supported on a 200-mesh copper grid (Plano GmbH, Germany) placed in the AES. The grids were analyzed using a Transmission Electron Microscope (JEM-2100F, 219 JEOL Ltd, Japan) operated at 200 kV, and images were acquired at 15,000x and 40,000x magnification.

### Cell model systems

The human lung adenocarcinoma, bronchial epithelial cell line Calu-3 (HTB-55, ATTC) was cultured in minimum essential medium (MEM)(1x) + GlutaMAX -I (41090-093, GIBCO) supplemented with 10% sterile filtered and heat inactivated fetal bovine serum (FBS) Honduras Origin Sterile Filtered (F7524-500ML, SIGMA), 1% Penicillin-Streptomycin (15140-122, GIBCO) and 1% MEM non-essential amino acids (NEAA) (1140-050, GIBCO). The human monocyte cell line THP-1 (ATTC, TIB-202), the human somatic cell hybrid endothelial cell line EA.hy926 (CRL-2922, ATTC,) and the human lung carcinoma, alveolar-epithelial cell line A549 (CCL-185, ATTC) were cultured in Dulbecco’s Modified Eagle Medium (DMEM)/F-12 (1:1) (1X) + GlutMAX-I (31331-093, GIBCO) supplemented with 10% FBS and 1% Penicillin-Streptomycin. After three passages A549 cells were set to 5% FBS. All cells were kept in a cell incubator in humidified air with 5% CO_2_ at 37 °C.

Monocytic THP-1 cells were differentiated into macrophages (dTHP-1) with cell culture medium supplemented with 250 nM Phorbol-12-myrstate-13-acetate (524400, SIGMA) reconstituted in DMSO, for 24 h, followed by 24 h resting time.

For assembly of the 3D model, on day zero 500,000 (equivalent to 107,000 cells/cm^2^) Calu-3 cells were seeded in submerged conditions on Transwell Permeable Supports, 24 mm, 6-well plate, 0.4 µm polyester membrane (3450, COSTAR). Every other day cells were washed with Hanks’ Balanced Salt Solution (HBSS) (14025-050, GBICO) and medium was renewed. On day seven cells were set to ALI. Calu-3 cells were cultured at ALI for 7 days to ensure optimal differentiation respectively maturation. On day 14 inserts were inverted and 100,000 EA.hy926 cells were seeded at the basolateral side of the membrane. After attachment of EA.hy926 cells, inserts were placed back in the plate, and 126,000 dTHP-1 cells were seeded on top of the Calu-3 cells. On day 15 the 3D model was used for exposure and experiments.

For the A549 model, on day zero 250,000 A549 cells were seeded on transwell inserts (54,000 cells/cm^2^). On day two cells were set at the ALI and the medium in the basolateral compartment was renewed. On day three the medium in the basolateral compartment was renewed. On day four the model was used for exposure experiments.

### Exposure of model systems at the air-liquid interface

Inserts seeded with the 3D or the A549 model were placed in the AES with 6.5 mL of exposure medium in the buckets of the AES constituting the basolateral compartment, as previously described by (Offer et al. 2022). Exposure medium consisted of minimum essential medium supplemented with 1% Penicillin-Streptomycin, 1% non-essential amino acids and 15 mM HEPES Buffer Solution (1M) (15630-056, GIBCO) for the 3D model or DMEM supplemented with 1% Penicillin-Streptomycin and 15 mM HEPES for the A549 model. At the apical compartment the models were exposed for 4h to SDG-UFP or catalytically cleaned compressed air (CA) at the ALI at continuous flow rates of either 20, 60 or 100 mL/min for the 3D model, or 100 mL/min for the A549 model. The inlet flow of the exposure station was set to 500 L per hour. The exposure station was heated to 37°C. Prior to exposure the aerosol and the CA were conditioned to 37°C and 85% relative humidity. Three independent biological replicates of each condition were conducted.

### Inductively coupled plasma mass spectrometry (ICP-MS) for SDG-UFP deposition measurement

Directly after exposure the insert membranes of the 3D model were cut out and frozen at -20 °C for storage. Membranes were subjected to microwave assisted pressurized acidic digestion and the mass of elemental copper was measured by a Triple Quadrupole inductively coupled plasma mass spectrometer (ICP-MS) (8900 QqQ, Agilent) as previously described (Cao et al. 2022, Neukirchen et al. 2024). By subtracting the mass of copper measured in CA exposed cells the mass deposition by 20, 60 and 100 mL/min flow rate SDG-UFP aerosol was determined. Results are given as average of three independent replicates as mass of deposited copper per surface area of the insert membrane (4.67 cm^2^). The average aerosol mass feed was calculated by multiplying the average copper concentration of the SDG-UFP aerosol (Das et al. 2025) with the aerosol feed from the AES by a flow rate of 20, 60 or 100 mL/min over an exposure duration of 4h.

### Lactate dehydrogenase (LDH) assay

After exposure, the apical compartment of the cell insert was washed with 1mL pre-warmed HBSS. Lactate dehydrogenase (LDH) release in the wash and the conditioned exposure medium from the basolateral compartment was quantified by using a Cytotoxicity Detection Kit (11644793001, Roche) according to manufacturer’s instructions. 100 µL of sample were measured in 96-well format. The colorimetric reaction was stopped after 7 min. LDH was quantified by measuring the absorbance at 493 nm with a multimode microplate reader (Varioskan LUX, Thermo Fischer Scientific). HBSS or exposure medium was used for blank correction. For determination of maximal LDH value incubator control cells were lysed with 2% Triton X-100 (BJ3924, Sigma-Aldrich) in HBSS for 15 min and measured in the same way. LDH release is given as the percentage of cytotoxicity as average of three independent experiments.

### Resazurin reduction assay

Metabolic activity was measured by the intracellular reduction of Resazurin to the highly fluorescent resorufin, commonly known as alamar-blue assay. After the exposure the apical compartment of the cell insert was washed with 1 mL pre-warmed HBSS. The HBSS was removed, the inserts were placed in new 6-well plate and 1 mL of 10% PrestoBlue Cell Viability Reagent (A13261, Invitrogen) in prewarmed cell culture medium was added to both compartments. Inserts were incubated in a cell incubator for 1 h and cell viability was accessed by measuring fluorescence with excitation at 493 nm and emission at 590 nm with a multimode microplate reader (Varioskan LUX, Thermo Fischer Scientific). Metabolic activity was accessed by subtracting a blank control (10% PrestoBlue reagent) and subsequent adding up of apical and basolateral fluorescence values. Results are given as fold change (FC) of metabolic activity over incubator control (IC) cells, as average of three independent experiments.

### Trans epithelial electrical resistance (TEER)

After exposure inserts were transferred to 6-well plates filled with 1mL of pre-warmed HBSS. 1mL of pre-warmed HBSS was added to the apical compartment and the inserts were incubated for 10 min in a cell incubator. Electrical resistance was measured with an Epithelial Volt/Ohm Meter 3 (EVOM3, World Precision Instruments). TEER was calculated by subtracting the resistance of a blank measurement and multiplying the resistance with the insert surface area. Results are given as the average TEER of three independent experiments.

### Transcriptome analysis

After exposure the apical compartment was washed with 1 mL pre-warmed HBSS, after removal of the wash 1mL of RNA protect cell reagent (76526, QIAGEN) was added to both compartments and the cells were harvested by scratching them of the inserts and frozen at -80 °C. RNA extraction from the frozen cells was performed with an RNA Plus Mini Kit (74136, QIAGEN) according to the manufacturer’s instructions. RNA sequencing (RNA-seq) analysis was performed at the Genomics Core Facility at Helmholtz Munich (Germany). After RNA isolation, RNA integrity number (RIN) was measured using an Agilent Fragment Analyzer 5300 system. RNAs with a RIN value > 7 were selected for totRNA sequencing (ribo-depleted). The libraries were prepared using the Illumina Stranded totalRNA Prep, Ligation kit with Ribo-Zero plus (Illumina), following the kit’s instructions. After a final quality check, the libraries were sequenced in a paired-end mode (2×100 bases) in the NovaseqX+ sequencer (Illumina) with a depth of ≥ 30 million paired-reads per sample. Only significant genes were considered (-log_10_ p-value ≥ 1.3) and a threshold of 1.5 FC over CA was applied (log_2_ fold-change ≥ 0.58 or ≤ -0.58). Subsequently, downstream analysis was conducted using the Ingenuity Pathway Analysis (IPA) software (QIAGEN). Pathways were selected according to the degree of activation or inhibition of biological processes predicted by a z-score ≥2 or ≤2, significance p-value (-log_10_ p-value ≥ 1.3) and biological relevance in the context of the study. Upstream regulators were identified by IPA software from the DEGs lists as described by (Kramer et al. 2014). Volcano plots to visualize gene expression data were generated with IPA software.

### Malondialdehyde measurement

After exposure, an aliquot of the conditioned exposure medium from the basolateral compartment was frozen at -80 °C for later measurement of Malondialdehyde (MDA). MDA was measured from the collected sample by a liquid chromatographic (LC) Triple Quadrupole MS/MS system (API 4000, AB Sciex), in positive MRM mode as previously described by (Cao et al. 2022). Results are presented as the average MDA concentration (ng/mL) from three independent experiments. Since the flow rate did not affect either MDA release or cytotoxicity in CA exposed 3D cultures, the MDA release from CA exposed cells is shown as the mean across all flow rates tested (20, 60 and 100 mL/min).

### Single cell gel electrophoresis (Comet assay)

After exposure, cells were washed with pre-warmed HBSS subsequently cells in both compartments were harvested with 0.025% Trypsin-EDTA Solution (T4174-100ML, Sigma). The cell suspensions were diluted, and the mini-gel version of the alkaline comet assay was performed as previously described (Di Bucchianico et al. 2017). As positive control an aliquot of incubator control cell suspension was incubated for 5 minutes with 30 µM H_2_O_2_ (Merk Milipore, 107209) at 4°C prior to gel preparation. Electrophoresis was carried out at 1.2V/cm^2^ at 270-300 mA for 25 min. Slides were stained with 1:10000 SYBR Gold Nucleic Acid Gel Stain (S11494, Invitrogen). Pictures of the mini gels were taken with the Lionheart FX automated microscope in 20x magnification. At least 100 comets per gel were scored with CometScore 2.0 (Rex Hoover Software, RexHoover.com, version 2.0.0.38). Results are given as the percentage of DNA in tail (% DNA in Tail) of three independent replicates.

### O-link/Proximity extension assay

After exposure samples from the basolateral medium were taken and frozen at -80 °C. Protein biomarker profiling of an “inflammation” panel and an “immune response” panel (92 molecules each) was conducted using O-link Proximity Extension Assay (PEA) technology based multiplex method (O-link proteomics, Thermo Fischer Scientific). The analysis was performed by the Core Facility Metabolomics and Proteomics, at Helmholtz Munich, Germany, a certified provider of O-link services. This method utilizes dual antibody binding and DNA barcoding to quantify protein biomarkers, generating Normalized Protein expression (NPX) values for downstream statistical analysis (Wik et al. 2021). The data was processed and quality-controlled according to the manufacturer’s guidelines and subsequently analyzed by IPA software (QIAGEN). For pathway analysis a p-value of overlap threshold of 0.15 (equivalent to a -log_10_ p-value ≥ 0.82) was chosen for the biomarkers.

### Enzyme-linked immunosorbent assay (ELISA)

Conditioned cell culture medium from the basolateral compartment and HBSS washes of the apical compartment were collected and frozen at -80 °C. After thawing up the concentration of the cytokines interleukin 6 (IL-6), IL-8 and IL-1β were measured with Human IL-6 DuoSet ELISA (DY206), Human IL-8/CXCL8 DuoSet ELISA (DY208) and Human IL-1 beta/IL1F2 DuoSet ELISA (DY201) (Bio-Techne) respectively. Assays were carried out in 96 plate scale with 100 µL sample volume according to manufacturer’s instructions. 3,30,5,50-tetramethylbenzidine (7004P6, Cell Signaling Technology) was used as chromogenic substrate and the absorbance was measured at 450nm and wavelength correction at 540 nm with a multimode microplate reader (Varioskan LUX, Thermo Fischer Scientific). Results are given as the mean cytokine concentration in pg/mL of three independent replicates in the basolateral medium for the A549 model and for the apical wash and the basolateral medium for the 3D model. Cytokine release from CA exposed 3D model is shown as mean across all flow rates tested (20, 60 and 100 mL/min).

### Statistical analysis

Unless otherwise stated, all statistical analysis were performed using GraphPad Prism 10 (version 10.6.0 (890), GraphPad Software). Statistical significance was determined using unpaired t-test or ordinary one-way ANOVA followed by Bonferroni’s multiple comparison test with a 95% confidence interval. Data are presented as the mean of at least three independent experiments, with errors bars indicating the standard error of the mean (SEM). Graphs were generated using GraphPad Prism 10.

## Results

### Aerosol characterization and particle deposition on the 3D and A549 model

TEM micrographs (Fig. 1a and 1b) revealed that the SDG-UFP generated by spark ablation were predominantly present as loosely agglomerated nanoscale primary particles with irregular, chain-like cluster morphologies and a relatively homogeneous spatial distribution across the grid, consistent with typical spark-generated aerosol structures. Their physical characteristics, including size distribution, particle number concentration, and geometric mobility diameter, were previously reported by Das et al. (2025) and are briefly summarized here. Spark ablation generated copper UFP with a mean mobility diameter of 33 ± 2 nm, with most particles below 100 nm (Fig. 1c). The average SDG-UFP aerosol mass concentration was 80 ± 10 µg/m^3^, and the mean particle number concentration (PNC) was 1.1×10^6^/cm^3^ ± 0.2×10^4^ (Das et al. 2025). For dose-response assessment, the 3D model was exposed for 4 h to continuous SDG-UFP flows of 20, 60, or 100 mL/min. ICP-MS analysis showed deposited copper surface masses of approximately 6.5, 21.8, and 41 ng/cm^2^, respectively (Fig. 1d). These values correspond to 7.9%, 8.8%, and 10% of the average aerosol mass introduced into the exposure system. Although deposition on the A549 model was not directly measured, similar behavior was assumed because exposure conditions were identical and only minor surface topography differences existed between models.

**Fig. 1:**
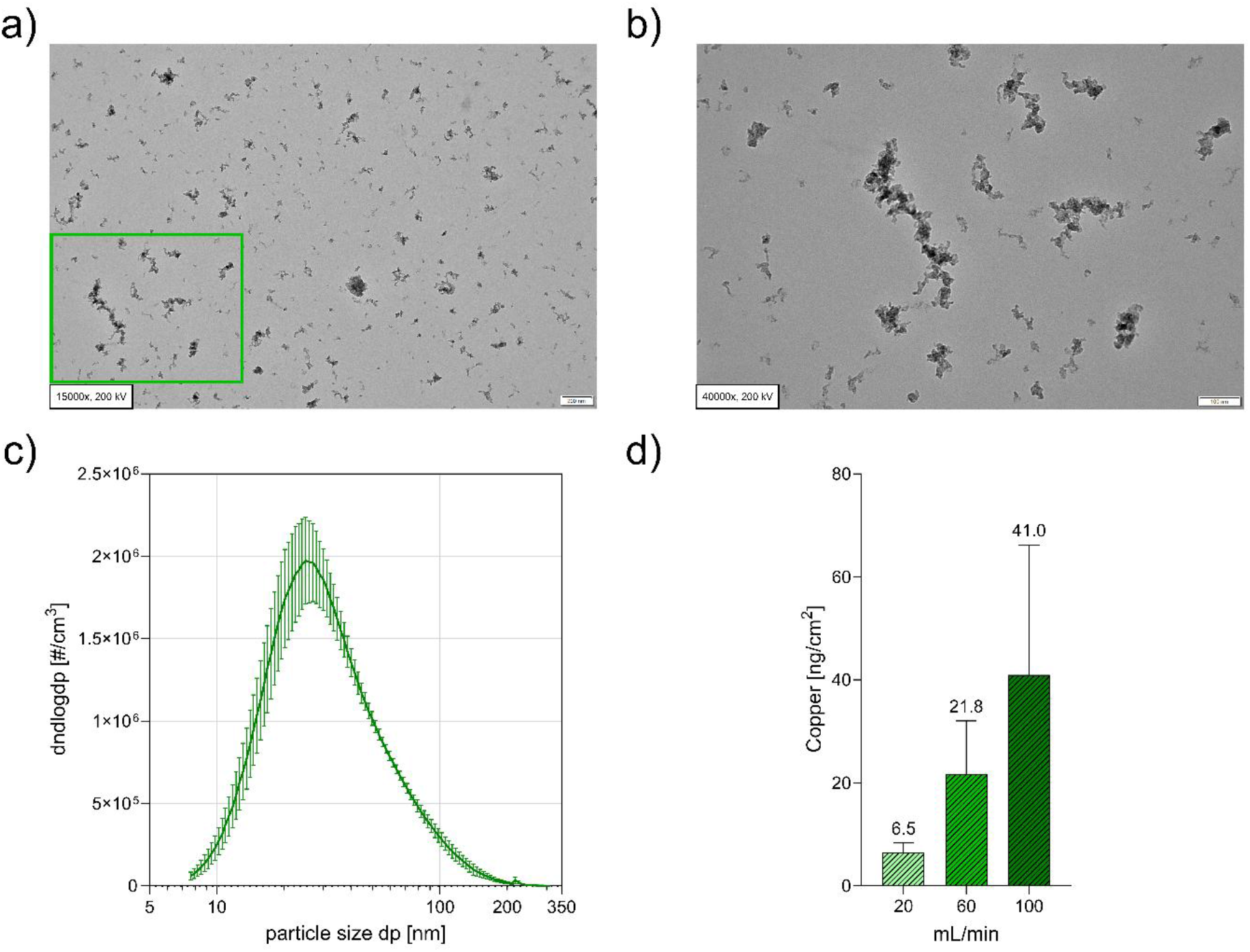
SDG-UFP physical characteristics and resulting mass deposition. TEM images of SDG-UFP, (**a**) overview at 15,000x and (**b**) 40,000x of the area indicated in green in (a). (**c**) SDG-UFP size distribution. Error bars indicate the standard deviation of the mean number concentration (#/cm^3^) for each size bin. (**d**) Deposited SDG-UFP mass per cell insert surface area measured by ICP-MS following ALI exposures at flow rates of 20 mL/min (light green), 60 mL/min (green), or 100 mL/min (dark green). Data are presented as mean + SEM (N=3). Numerical values are indicated above the bars

### Biological response of two lung cell models to SDG-UFP

To assess acute cytotoxicity of SDG-UFP, LDH release was measured in both A549 cells and the 3D model. In A549 cells, exposure significantly increased cytotoxicity to 13.3%, indicating pronounced cell membrane damage (Fig. 2a). In contrast, the 3D co-culture model showed only modest LDH release across all flow rates (20, 60, and 100 mL/min), ranging from 6.4% to 7.8% (Fig. 2b). Cell viability, determined by resazurin reduction, was slightly reduced after exposure at 60 and 100 mL/min to 91.2% and 91.7%, respectively (Fig. 2c). Barrier integrity, assessed by TEER, was significantly impaired at the same flow rates, decreasing to 624.5 Ω·cm^2^ and 507.4 Ω·cm^2^, respectively (Fig. 2d). Overall, A549 monocultures were more susceptible to SDG-UFP-induced membrane damage, whereas the 3D co-culture model largely preserved viability despite marked disruption of epithelial barrier function.

**Fig. 2:**
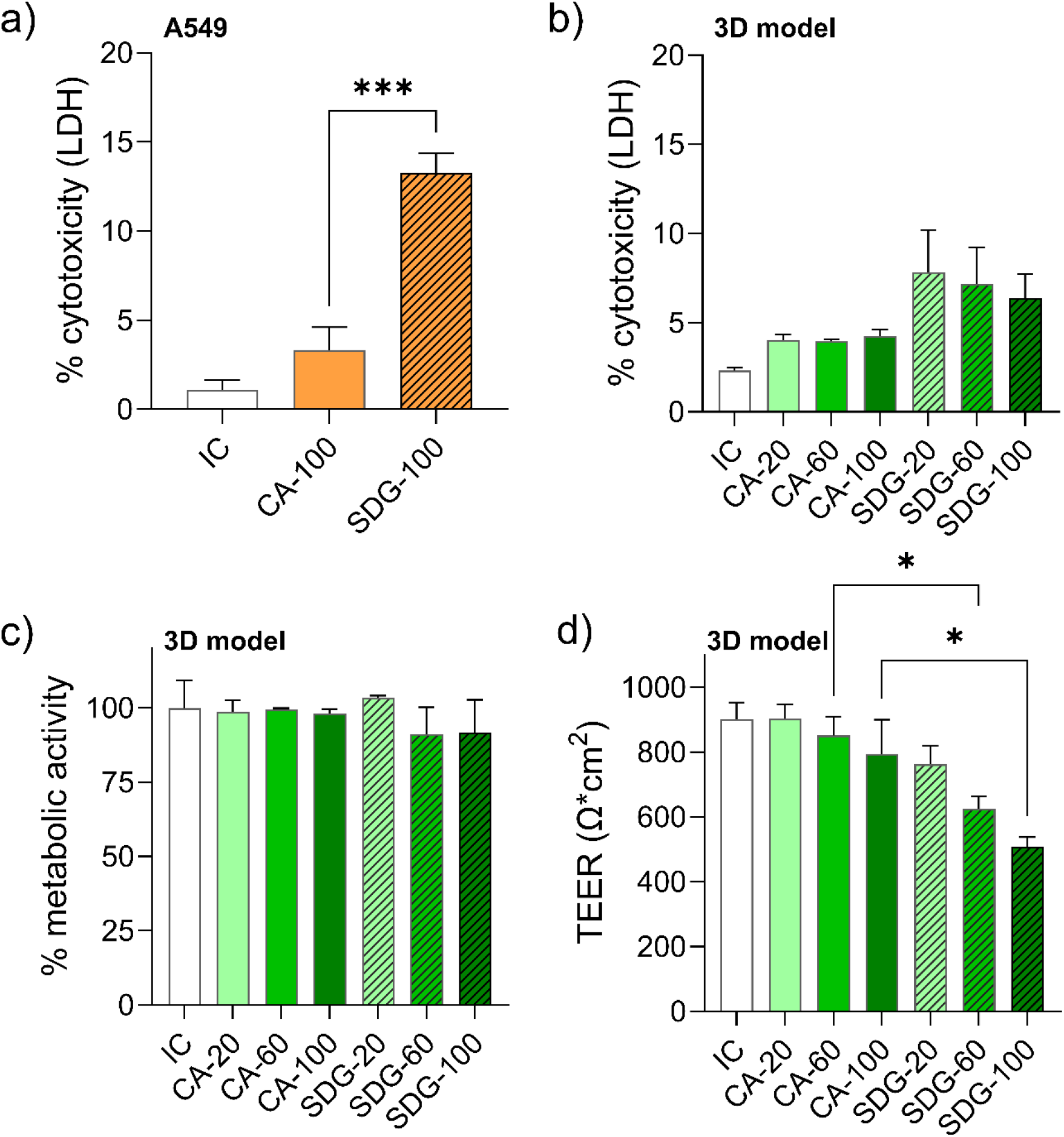
Cellular health of A549 cells (orange) and 3D model (green). (**a**) Cytotoxicity (%), measured by lactate dehydrogenase (LDH) release, in A549 cells exposed to 100 mL/min clean air (CA, orange) or 100 mL/min SDG-UFP (orange, dashed). (**b**) Cytotoxicity (%) in the 3D model exposed to 20, 60 and 100 mL/min CA (green) or SDG-UFP (green, dashed). (**c**) Metabolic activity (%) in the 3D model under the same exposure conditions. (**d**) Trans epithelial electrical resistance (TEER) in the 3D model under the same exposure conditions. White bars indicate incubator control (IC). Data are presented as mean + SEM (N=3), * p-value ≤ 0.05, *** p-value ≤ 0.005

### Transcriptional changes in SDG-UFP exposed cells

Given that SDG-UFP exposure induced minimal cytotoxicity in the 3D and only modest cytotoxic effects in the A549 model, we performed transcriptomic profiling via RNA-seq of cells exposed to the highest flow rate (100 mL/min) to obtain sensitive and mechanistic insights into the molecular basis of SDG-UFP toxicity. The compartmentalized architecture of the 3D model enabled independent harvesting and analysis of Calu-3 + dTHP-1 and endothelial EA.hy926 cells.

In the A549 model, 308 differentially expressed genes (DEGs) were identified relative to control-exposed cells (-log_10_ p-value ≥ 1.3; log_2_ fold-change ≥ 0.58 or ≤ -0.58), with the majority (242 DEGs; 78.6%) showing upregulation (supplementary material, Fig. S1). In contrast, the 3D model exhibited a more balanced transcriptomic response, with 129 of 232 DEGs (55.6%) upregulated in Calu-3 and dTHP-1 cells and 61 of 138 DEGs (44.2%) upregulated in endothelial cells (supplementary material, Fig. S2 and S3). Cross-model comparison revealed limited transcriptomic concordance: only 14 DEGs overlapped between A549 and Calu-3 + dTHP-1 cells, four DEGs between A549 and endothelial cells, and three DEGs were shared between Calu-3 + dTHP-1 and endothelial compartments, highlighting the cell type-specific nature of particle-induced molecular responses (Fig. 3a).

**Fig. 3:**
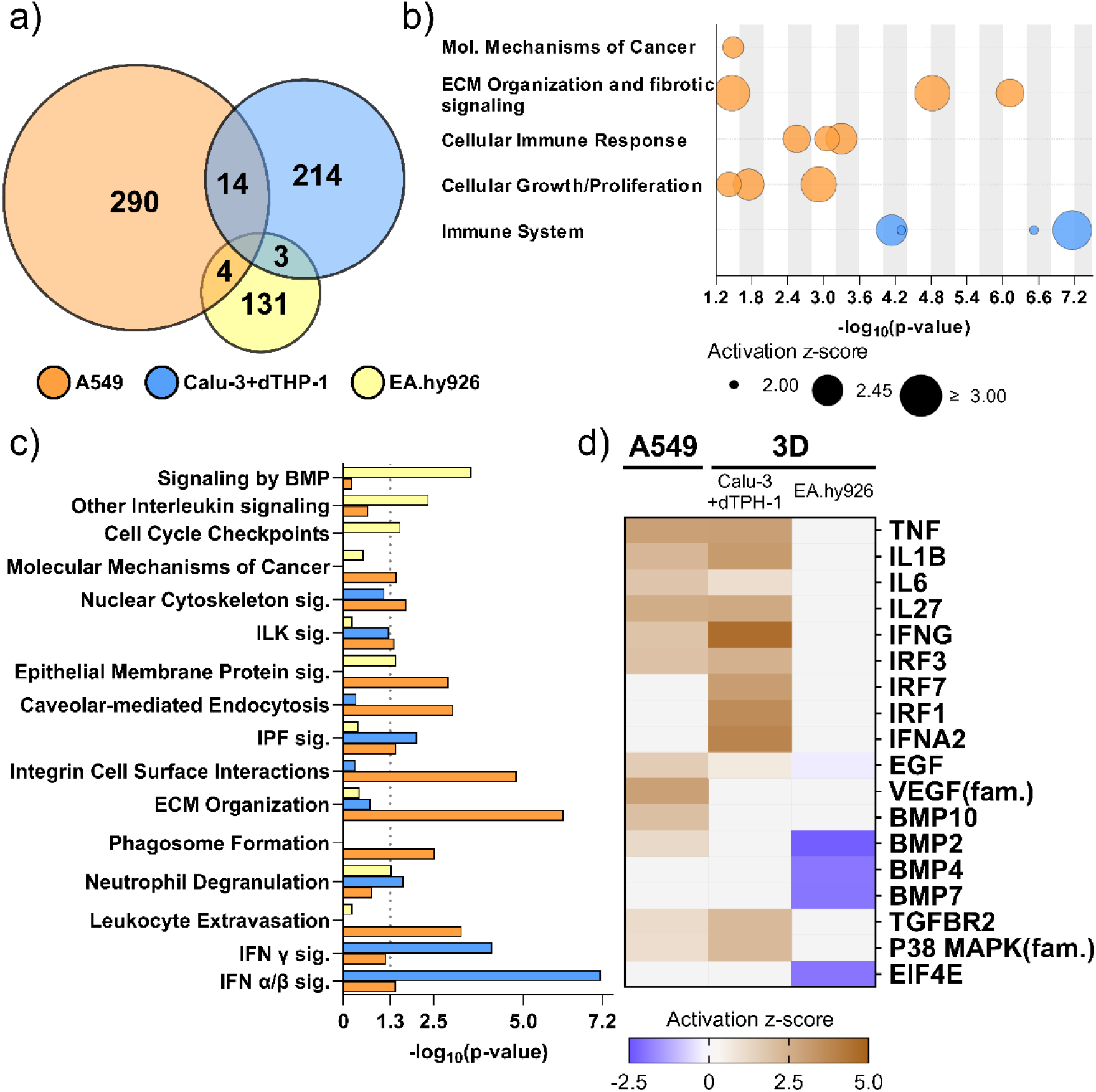
Transcriptome analysis of RNA-seq data of A549 (orange), Calu-3 + dTHP-1 (blue) and EA.hy926 (yellow) cells exposed to SDG-UFP at 100 mL/min. (**a**) Venn diagram of the overlapping and unique DEGs (-log_10_ Expr. p-value ≥1.3, log_2_ Expr. FC ≥ 0.58 and log_2_ Expr. FC ≤ -0.58). (**b**) Clustering of relevant, activated canonical signaling pathways (bubbles) to biological processes (y-axis). Bubble size represents z-score as measure for pathway activation. (**c**) Results of the ingenuity pathway analysis of the DEGs in (a) showing the significance of overlap (y-axis) for a selection of relevant pathways (-log_10_ p-value of overlap ≥ 1.3). (**d**) Activation of a selection of relevant upstream regulators governing the pathways depicted in B and C (-log_10_ Expr. p-value ≥ 1.3). All data represents mean of three independent experiments

Mapping of DEGs to the IPA canonical pathway database revealed associations with 106 signaling pathways in A549 cells, 42 pathways in Calu-3 + dTHP-1 cells, and 49 pathways in endothelial cells (-log_10_ p-value ≥ 1.3). Comprehensive pathway analysis integrating activation z-scores, biological context, and cell type-specific considerations identified distinct SDG-UFP-induced biological processes in directly exposed cells (Fig. 3b). In A549 cells, SDG-UFP exposure activated pathways governing extracellular matrix (ECM) organization, profibrotic signaling, cellular growth and proliferation and cellular immune responses, with additional associations to molecular mechanisms of cancer. The Calu-3 + dTHP-1 compartment exhibited dominant immune system activation. In the endothelial EA.hy926 cells the DEG pattern significantly overlapped with several pathways concerning growth and immune signaling however, no prediction of activity (z-score) was possible due to the insufficient number of DEGs.

Dysregulation of ECM organization and profibrotic processes in A549 cells was characterized by activation of the idiopathic pulmonary fibrosis (IPF), integrin cell surface interactions, and ECM organization pathways (Fig. 3c). This molecular signature was substantiated by enhanced expression of multiple ECM-associated genes, including integrin-α1 (*ITGA1*, log_2_ fold-change = 0.76, -log_10_ p-value = 1.55), integrin-α2 (*ITGA2*, log_2_ fold-change = 1.05, -log_10_ p-value = 3.21), integrin-β4 (*ITGB4*, log_2_ fold-change = 0.64, -log_10_ p-value = 2.40), and integrin-β8 (*ITGB8*, log_2_ fold-change = 0.59, -log_10_ p-value = 1.97), as well as fibronectin-1 (*FN1*, log_2_ fold-change = 0.72, -log_10_ p-value = 2.01), heparan sulfate proteoglycan-2 (*HSPG2*, log_2_ fold-change = 0.67, -log_10_ p-value = 2.17), and collagen IV α-4 (*COL4A4*, log_2_ fold-change = 0.69, -log_10_ p-value = 2.33). Concurrently, key intracellular signal-transducing kinases mitogen-activated protein kinase 12 (*MAPK12*, log_2_ fold-change = 0.64, -log_10_ p-value = 1.52) and protein tyrosine kinase 2 beta (*PTK2B*, log_2_ fold-change = 1.08 -log_10_ p-value = 2.80) exhibited significant upregulation. These genes additionally contributed to activation of the epithelial membrane protein and nuclear cytoskeleton signaling, as well as integrin-linked kinase (ILK) pathways, which collectively govern cellular growth and proliferation processes. Moreover, we found activation of the molecular mechanisms of cancer pathway. Cellular immune responses were activated through increased leukocyte extravasation signaling, phagosome formation, and caveolar-mediated endocytosis pathways (Fig. 3b and 3c). Calu-3 + dTHP-1 cells demonstrated robust immune system activation, with pathway analysis revealing significant upregulation of canonical interferon (IFN) α/β signaling. This immune response was reflected by elevated expression of key IFN-stimulated genes, including IFN-induced protein with tetratricopeptide repeats (*IFIT*1, log_2_ fold-change = 1.27, -log_10_ p-value = 1.64), *IFIT2* (log_2_ fold-change = 1.92, -log_10_ p-value = 1.90), and *IFIT3* (log_2_ fold-change = 1.58, -log_10_ p-value = 1.63), as well as IFN alpha and beta receptor subunit 2 (*IFNAR2*, log_2_ fold-change = 0.67, -log_10_ p-value = 1.64). Notably, these genes were also annotated to neutrophil degranulation processes. Additionally, transcripts encoding secreted mucins *MUC1* (log_2_ fold-change = 1.37, -log_10_ p-value = 1.98) and *MUC5B* (log_2_ fold-change = 1.41, -log_10_ p-value = 1.88) were significantly elevated in the apical compartment. In endothelial EA.hy926 cells highest significance was found for bone morphogenetic protein (BMP) and other interleukin signaling pathway (Fig. 3c). Moreover, the cell cycle checkpoints pathway was found downregulated (-log_10_ p-value = 1.59, z-score = -1.00).

Upstream regulator analysis was performed to identify molecular entities activated upstream of the DEGs, thereby elucidating the root causes of the observed transcriptomic alterations (Fig. 3d). Both models (A549 and Calu-3 + dTHP-1) demonstrated activation of secretory cytokines including tumor necrosis factor (*TNF*), interleukin-1 beta (*IL1β*), *IL-6*, and *IL-27*. The 3D model exhibited pronounced activation of the interferon network, with IFN-gamma (*IFN-γ*) identified as the highest-ranking activated upstream regulator, followed by IFN-alpha 2 (*IFNA2*), IFN regulatory factor 1 (*IRF1*), and *IRF7*. Conversely, the A549 model showed pronounced activation of growth factor signaling, including epidermal growth factor (*EGF*), the family of vascular endothelial growth factor (*VEGF*), as well as *BMP10*. Both models demonstrated activation of the transforming growth factor beta receptor 2 (*TGFBR2*) and the signal transducing p38 mitogen-activated protein kinase (MAPK) family. Uniquely, the basolateral endothelial compartment in the three-dimensional model displayed exclusive inhibition of *BMP2, BMP4*, and *BMP7*, alongside inhibition of eukaryotic translation initiation factor 4E (*EIF4E*) (Fig. 3d).

In summary, SDG-UFP exposure elicited distinct cell type-specific transcriptomic responses across the respiratory models. A549 epithelial cells exhibited prominent activation of growth factor signaling and modest inflammatory responses and ECM remodeling and fibrosis signaling. The apical cells of the 3D model (Calu-3 + dTHP-1) demonstrated dominant interferon-mediated immune activation, reflecting enhanced innate immune defense mechanisms. The basolateral endothelial compartment (EA.hy926) was characterized by inhibition of bone morphogenetic protein signaling and reduced activation of the cell cycle pathway.

### Induction of oxidative stress and genotoxicity of SDG-UFP

MDA concentration, a marker of oxidative stress, was measured in the basolateral medium. SDG-UFP exposure had no effect on MDA concentration in the A549 model (Fig. 4a). In contrast, exposure in the 3D model resulted in a significant increase in MDA at 100 mL/min, while exposure at 20 and 60 mL/min induced modest but non-significant increases (Fig. 4b). Genotoxicity was assessed using the alkaline comet assay, which quantifies alkali-labile DNA lesions and strand breaks. Cells were exposed at the highest flow rate of 100 mL/min. No DNA damage was observed following exposure of A549 (Fig. 4c). In the apical compartment of the 3D model, a slight increase in DNA strand breaks was detected (Fig. 4d), however, no DNA damage was observed in the underlying endothelial cells (Fig. 4e). Collectively, these findings indicate elevation of oxidative stress markers in the basolateral compartment accompanied by concurrent emergence of genotoxic effects in directly exposed apical cells of the 3D model.

**Fig. 4:**
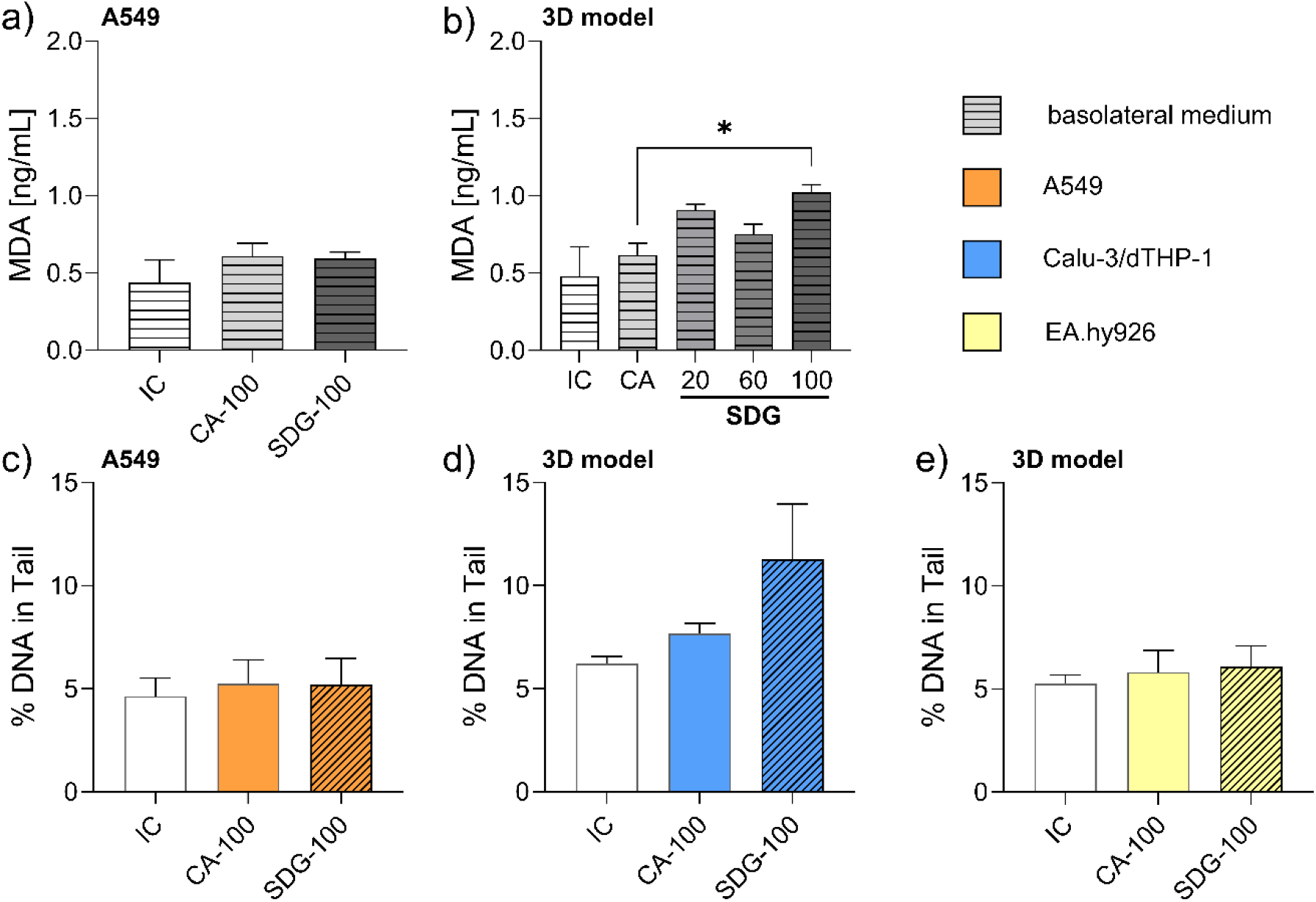
Oxidative stress and DNA damage in the SDG-UFP exposed A549 and 3D model. (**a** and **b**) Concentration of the oxidative stress marker Malondialdehyde (MDA) in the basolateral medium (dashed) of the A549 model (**a**) and the 3D model (**b**) exposed to clean air (CA) or SDG-UFP at different flow rates (mL/min). (**c**-**e**) DNA damage, measured by percent DNA in tail in the alkaline comet assay in the A549 model (c, orange), the apical compartment of the 3D model (d, blue) and the basolateral compartment of the 3D model (e, yellow) exposed to CA or SDG-UFP (dashed) at a flow rate of 100 mL/min. White bars indicate incubator control (IC) cells, all data shown as mean + SEM (N=3), * p-value ≤ 0.05

### Proteome analysis of the basolateral exposure medium

Proteome analysis of basolateral medium demonstrated enrichment of selected biomarkers in the 3D model whereas the A549 model showed predominantly depleted responses. Among the 184 markers assessed, the 3D model exposed to flow rates of 20, 60, and 100 mL/min yielded a total of 25, 35, and 22 significant hits, respectively, whereas the A549 model exposed to 100 mL/min SDG-UFP demonstrated 24 significant hits (−log_10_ p-value ≥ 0.82) (Fig. 5a). Twelve biomarkers displayed differential expression between the two models; in the 3D model their levels increased with exposure concentration, whereas only two biomarkers were upregulated in the A549 model (Fig. 5b). This included chemokine (C-C motif) ligand (CCL)2, CCL3, CCL4, IFN-γ, IL-6, IL-8, and chemokine (C-X-C motif) ligand (CXCL)1 which were predominantly elevated in 3D model conditions. At the highest exposure concentration, the 3D model additionally secreted CXCL5, matrix metalloproteinase-1 (MMP-1), tumor necrosis factor receptor superfamily member 9 (TNFRSF9), TNFRSF11B, and amphiregulin (AREG). Notably, IFN-γ was uniquely enriched in the 3D model while being significantly suppressed in the A549 model (Fig. 5b). Pathway enrichment analysis mapped these biomarkers to signaling cascades associated with chronic obstructive pulmonary disease (COPD) and cytokine storming, as well as IL-10 and IL-17 pathways.

**Fig. 5:**
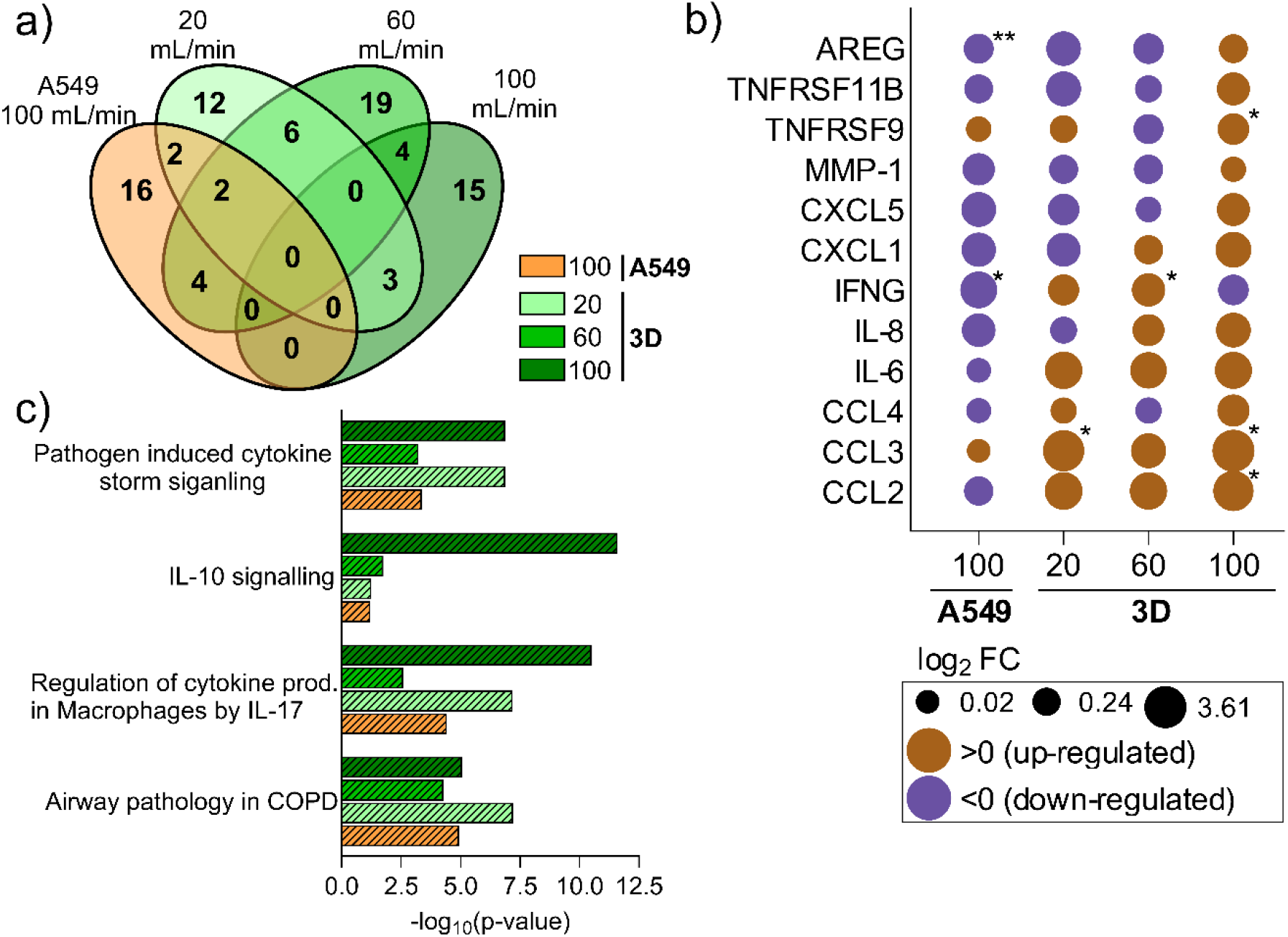
Proteome analysis biomarkers of inflammation in the conditioned medium of the A549 (orange) and 3D model (green). (**a**) Venn diagram of the overlapping and unique markers (-log_10_ p-value ≥ 0.82). (**b**) Enrichment (brown) or depletion (purple) of selected inflammatory markers across the 3D and the A549 model. Size represents log_2_ fold change (FC) over clean air (CA), * -log_10_ p-value ≥ 1.3, **-log_10_ p-value ≥ 1.99. (**c**) Overlap of molecules from (a) with relevant canonical signaling pathways indicated by -log_10_ p-value of overlap (x-axis). All data shown as mean of three independent experiments

### Secretion of inflammatory cytokines by the lung models

RNA-seq and proteomic analyses (Fig. 3 and 5) indicated robust inflammatory activation; therefore, pro-inflammatory cytokines (IL-1β, IL-6, and IL-8) were quantified by ELISA to functionally validate these findings.

In the A549 model, IL-8 was the only cytokine detected above the limit of quantification (LOQ), and its concentrations did not differ significantly between SDG-UFP and control conditions. Notably, the basolateral IL-8 concentration (25 pg/mL) was approximately twofold higher than the apical concentration (15 pg/mL) in SDG-UFP exposed cells (Fig. 6a and 6b).

**Fig. 6:**
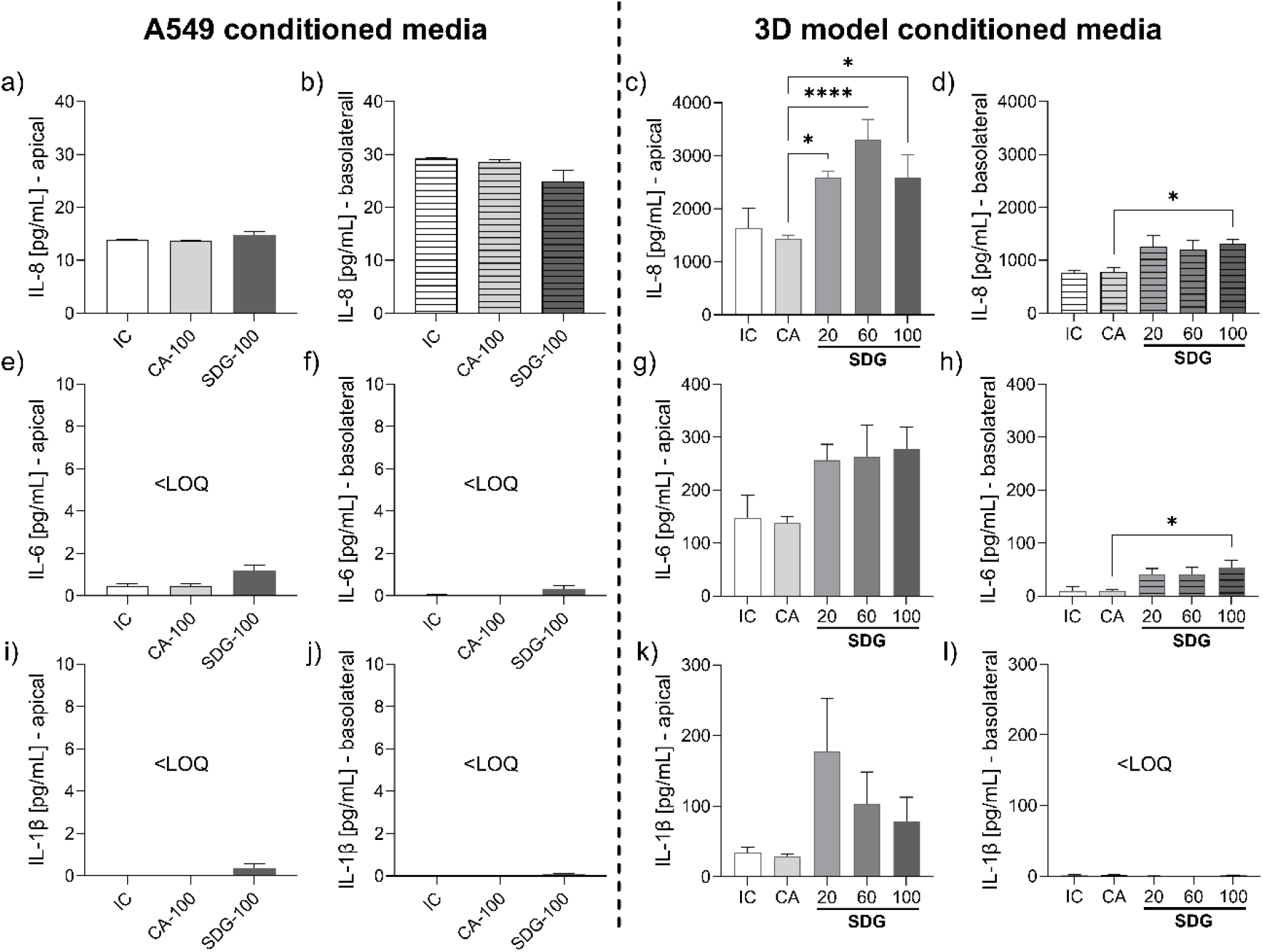
Quantification of pro-inflammatory cytokines in the conditioned medium of the A549 and 3D model by ELISA. (**a-d**) Concentration of interleukin (IL) IL-8 in the apical washes (a and c) and basolateral conditioned medium (b and d, dashed) of the A549 model (a and b) and 3D model (c and d) exposed to clean air (CA, light grey), SDG-UFP (dark grey) and in incubator control (IC, white). (**e**-**h**) Concentration of IL-6 in the apical washes (E+G) and basolateral conditioned medium (F+H, dashed) of the A549 model (e and f) and 3D model (g and h) exposed to CA, SDG-UFP and in IC. (**i**-**l**) Concentration of IL-1β in the apical washes (I+K) and basolateral conditioned medium (j and l, dashed) of the A549 model (I and j) and 3D model (k and l) exposed to CA, SDG-UFP and in IC. All data shown as mean + SEM (N=3), * p-value ≤ 0.05, ** p-value ≤ 0.01, **** p-value ≤ 0.001. LOQ (below limit of quantification)

In the 3D model, SDG-UFP significantly increases IL-8 release at the highest exposure concentration (100 mL/min) in both the apical and basolateral compartments (Fig. 6c and 6d). In the apical compartment, exposure concentrations of 20 and 60 mL/min also significantly elevated IL-8 levels (Fig. 6c). IL-6 levels were similarly increased at all exposure concentrations in the apical compartment (Fig. 6g), whereas the higher deposition levels significantly increased basolateral IL-6 release (Fig. 6h). Apical IL-1β concentrations were elevated under all exposure conditions (Fig. 6k), although with high inter-assay variability, while IL-1β remained below the LOQ in the basolateral compartment (Fig. 6l).

Overall, the 3D model exhibited markedly enhanced cytokine secretion in response to SDG-UFP exposure, with consistently higher concentrations in the apical compartment (Fig. 6c, 6g and 6k) compared to the basolateral (Fig. 6d, 6h and 6l). This pattern likely reflects the contribution of dTHP-1 cells localized to the apical compartment, as well as apically polarized secretion by Calu-3 cells, which interface with both compartments. The significantly increased IL-6 and IL-8 secretion in the 3D model corroborates inflammatory activation in line with the transcriptome profiling and proteomic findings, whereas IL-1β responses remain inconclusive due to high variability, despite a modest apical increase.

## Discussion

SDG-UFP deposition induced significant cytotoxic effects in the A549 model (Fig. 2a), consistent with prior observations on copper nanoparticles and highlighting the health hazard posed by ultrafine particles (Kwon et al. 2020).

At the RNA level SDG-UFP exposure triggered a coordinated dysregulation of ECM remodeling and activation of cellular growth and proliferation in A549 cells (Fig. 3b and 3c). Key ECM-associated genes, including *COL4A4* and *FN1*, were substantially upregulated, as were downstream signaling kinases *MAPK12* and *PTK2B*. Moreover, dysregulation of growth and proliferation processes was further reflected by activation of growth factors in the upstream regulator analysis (Fig. 3d). The concomitant activation of the idiopathic pulmonary fibrosis pathway is particularly noteworthy, as it mechanistically connects ECM dysregulation and proliferation signaling to pro-fibrotic programming (Fig. 3c) (Herrera et al. 2018, Zhou et al. 2024). Both *MAPK12* and *PTK2B* are known drivers of fibrotic responses through fibroblast activation and collagen synthesis promotion (Zhou et al. 2024). *In vivo* evidence demonstrates that copper oxide nanoparticle deposition in the murine lung leads to epithelial injury that can progress to fibrosis, characterized by elevated collagen production (Lai et al. 2018). Together, these data suggest that SDG-UFP exposure can initiate molecular pathways associated with chronic pulmonary fibrotic disease even at below genotoxic and ROS inducing concentrations (Fig. 4a and 4c).

A striking disconnect emerged between transcriptional and secreted inflammatory responses in the A549 model. Although RNA-seq revealed activation of cellular immune response (Fig. 3b) the conditioned medium and apical washes contained minimal detectable inflammatory biomarkers (Fig. 5 and 6). This is illustrated by the fact that although *IFN-γ* was identified as an active upstream regulator in the transcriptome (Fig. 3d) proteome analysis of the conditioned medium found its significant depletion (Fig. 5b). However, activation of phagosome formation and caveolar-mediated endocytosis signaling pathways may benefit particle internalization (Fig. 3c). Previously, copper nanoparticles have been detected within endocytic vesicles of A549 cells and other organelles (Wang et al. 2012). Translocation of inhaled UFP through the lung epithelium, by paracellular routes (Muhlfeld et al. 2008) is considered as a driver for secondary toxicity of inhaled particles and has been demonstrated for model nanoparticles in humans (Miller et al. 2017).

The absence of robust inflammatory activation despite transcriptional evidence of immune pathway engagement highlights a fundamental limitation of the A549 monoculture in this study especially considering the low dosimetry applied in ALI nanoparticle toxicity assessment (Paur et al. 2011). Complex particle-induced toxicity inherently involves the coordinated response of multiple tissue compartments, which co-culture represent more adequately (Rothen-Rutishauser et al. 2023). This is exemplified by comparative studies demonstrating that combined cultivation of A549 epithelial cells with differentiated macrophage-like cells (dTHP-1) elicits substantially enhanced pro-inflammatory responses to copper nanoparticles compared to epithelial monocultures alone (Hufnagel et al. 2021).

While SDG-UFP exposure elicited negligible effects on cytotoxicity and viability in the 3D model (Fig. 2b and 2c), comprehensive RNA-seq analysis revealed robust activation of innate immune responses, with interferon innate immune response signaling as the dominant regulatory axis (Fig. 3b and 3c). This was corroborated by upstream regulator analysis, which identified *IRF7* as a central molecular driver of this response (Fig. 3d). Consistently occupational field studies demonstrated that metal-rich particulate matter from welding processes triggers type I interferon responses *in vivo*, with *IRF7* serving as a central transcriptional regulator during sustained exposure (Erdely et al. 2012). *IFN-γ* and *IL-6* emerged as a critical active upstream regulator in the transcriptome (Fig. 3d), substantiated by proteomic analysis, which identified concentration-dependent elevations in IL-6, IL-8, CCL2, CCL3, and IFN-γ consistent with prior studies on lung toxicity of copper oxide nanoparticles (Fig. 5B) (Jing et al. 2015, Lai et al. 2018, Parkin et al. 2025). Notably, pathway analysis revealed that this inflammatory signature shares similarities with COPD and pathogen-induced cytokine storming (Fig. 5c), reflecting a pattern where IFN-γ mediated priming of bronchial epithelial cells amplifies IL-6 production and triggers cytokine-mediated tissue damage (Karki and Kanneganti 2021, Okuma et al. 2023).

Proteome findings were functionally validated by ELISA, confirming enhanced secretion of IL-6, IL-8, and IL-1β (Fig. 6). Mechanistically, IL-6 can modulate mucin production while IL-8 eventually downregulates tight junction proteins, thereby compromising epithelial barrier integrity (Chen et al. 2003, Yu et al. 2013). Consistent with these mechanisms, we observed increased *MUC1* and *MUC5B* mRNA expression accompanied by concentration-dependent impairment of TEER (Fig. 2d), paralleling observations in previous studies of copper nanoparticle-exposed Calu-3 and alveolar type II cells (Bengalli et al. 2013). The relatively modest cytotoxicity may reflect protective effects conferred by co-cultured macrophages and endothelial cells and mediated by secretory cytokines (Liu 2025), especially when considering the relevant IL-6 and IL-8 levels present when other endpoint were accessed.

Production of ROS increased at the highest exposure concentration as indicated by elevated MDA levels in the basolateral medium (Fig. 4b). Copper nanoparticles internalized in lysosomes are suspected to give rise to ROS by dissolution in this acidic environment (Karlsson et al. 2013). Interestingly, IL-6 confers protection of lung epithelial cells against oxidative stress, potentially explaining the smaller MDA production at low exposure concentrations (Fig. 4b) (Kida et al. 2005). Additionally, genotoxicity assessment revealed elevated DNA damage in Calu-3 and dTHP-1 cells at a deposited dose of 41 ng/cm^2^ (Fig. 4d). These findings demonstrate that SDG-UFP recapitulate the pro-inflammatory, genotoxic and oxidative stress inducing profile previously documented for copper nanoparticles (Hufnagel et al. 2020, Kuhn et al. 2025, Siivola et al. 2020).

We hypothesize that differentiated THP-1 cells served as the central effectors initiating the innate immune response, as ELISA revealed strong polarization of cytokine secretion toward the apical compartment (Fig. 6), and macrophage lung epithelial cell interactions have been shown to be critical for nanoparticle-induced neutrophilia *in vivo* (Liu et al. 2025). Pathway analysis further implicated dysregulation of macrophage-derived cytokine production (Fig. 5c). However, given that Calu-3 cells produce cytokines upon macrophage-mediated activation, they likely contribute substantially to the overall inflammatory secretion profile, especially in the basolateral conditioned medium, once activated (Braakhuis et al. 2020, Fr et al. 2021).

Overlap with other interleukin signaling and BMP signaling pathways (Fig. 3c) suggest that endothelial cells actively participate in sensing inflammatory signals. This is consistent with previous findings demonstrating that air-blood barrier models facilitate intercellular communication through cytokine exchange in response to nanoparticle exposure (Bengalli et al. 2013). However, the absence of broader inflammatory pathway activation in endothelial cells supports the hypothesis that inflammation signal production is predominantly mediated by Calu-3 and dTHP-1 cells in the apical compartment. Upstream regulator analysis (Fig. 3d) provided mechanistic insights into the transcriptomic alterations in endothelial cells. The deactivation of BMPs potentially reflects dysregulation of vascular integrity and angiogenic processes, as *BMP7* inhibition has been shown to suppress endothelial quiescence (Chen et al. 2018, Ricard et al. 2021). Conversely, *EIF4E* inhibition suggests a generalized suppression of protein synthesis, which is further supported by the concurrent downregulation of the cell cycle checkpoint pathway (Fig. 3c). Collectively, these findings indicate a multifaceted transcriptomic profile characterized by reduced vascular integrity and cell proliferation in endothelial cells. These observations are noteworthy given that particulate matter exposure represents a significant risk factor for cardiovascular disease, potentially inducing substantial alterations in vascular remodeling and wound healing processes (Sagheer et al. 2024). Indeed, epidemiological evidence demonstrates that prolonged UFP exposure is associated with substantially elevated cardiovascular morbidity and mortality (Downward et al. 2018). The exposure concentration used in this study (80 µg/m^3^) falls within the range of ambient PM_2.5_ values reported for polluted subway systems (Loxham and Nieuwenhuijsen 2019). Data on UFP mass concentrations in such environments remain limited, however, PM_1_ concentrations of 21 µg/m^3^ and 37 µg/m^3^ have been reported in previous studies (Bendl et al. 2023, Loxham et al. 2013). Ambient air in these settings consists of mixed metal- and carbon-based particles, with copper representing only a fraction of total PM (Kumar et al. 2023). In contrast, occupational exposure scenario, such as industrial welding, can result in substantially higher copper concentrations. Accordingly, the United States National Institute for Occupational Safety and Health (NIOSH) has set an exposure limit of 100 µg/m^3^ for copper fumes (Brand et al. 2013, NIOSH 2007). Scientific bodies, including the European Commission’s Scientific Committee on Occupational Exposure Limits (SCOEL), have proposed a more conservative exposure limit of 10 µg/m^3^ for the respirable fraction of copper (SCOEL 2014).

The higher deposited SDG-UFP mass observed in this study (≈ 40 ng/cm^2^) corresponds to an estimated deposition efficiency of approximately 10%, based on aerosol concentration of 80 µg/m^3^. This value lies at the lower end of reported alveolar retention fractions (10-70%) from modelling and field studies (Paur et al. 2011, Su et al. 2024). Similarly, whole-lung deposition fractions for inhaled UFPs ranging from 28% to 70% have been reported in the literature (Daigle et al. 2003, Londahl et al. 2009). Based on modelling approaches, Paur and colleagues estimated a daily deposited dose ranging from 0.12 ng/cm^2^ under ambient conditions (10 µg/m^3^) to 130 ng/cm^2^ under worst-case occupational exposure scenarios, as a framework for realistic *in vitro* nanoparticle dosimetry (Paur et al. 2011). Accordingly, although the exposure concentration used here likely exceeds ambient UFPs levels in hotspots such as subway environments, the resulting deposited fraction in the AES is consistent with physiologically relevant lung deposition estimates from computational models (Karg et al. 2020).

Overall, the deposited range of doses fall within the levels encountered in highly polluted occupational settings, providing insight into the toxicity of copper-based UFPs. In addition, ALI exposure minimizes interactions between copper particles and culture medium components, enabling a more direct assessment of the intrinsic toxicity of UFPs generated by catenary sparking (Bessa et al. 2021). It should be noted that direct comparison of particle deposition between this *in vitro* system and ambient exposure scenarios is challenging, as differences in size distribution and aerosol density strongly influence deposition behavior and dosimetry (Grabinski et al. 2015).

## Conclusion

Subway systems are considered hot spots for metal PM exposure however only limited evidence exists on the UFP burden in such polluted environments. This study demonstrates that copper-based UFPs induce a complex, cell type-dependent response across lung-relevant *in vitro* models. In A549 monocultures, exposure resulted in clear cytotoxicity and transcriptional reprogramming consistent with extracellular matrix remodeling, proliferative signaling, and activation of pro-fibrotic pathways, suggesting early molecular events linked to chronic lung disease. In contrast, the 3D co-culture model exhibited minimal cytotoxicity but a pronounced innate immune response characterized by interferon-driven signaling, cytokine release, barrier dysfunction, and oxidative stress, more closely reflecting in vivo-like inflammatory processes.

Despite comparable or only modest deposited doses, ALI exposure to copper-based UFPs recapitulated key features of copper nanoparticle toxicity, including inflammation, genotoxicity, and oxidative stress, while revealing pronounced model-dependent differences in sensitivity. These findings further underscore the importance of multicellular ALI models in capturing intercellular communication and immune modulation that are absent in monocultures. Moreover, this study demonstrates for the first time the use of different airflow conditions within the same exposure setup, in both clean air and aerosol exposure modules, to assess aerosol dose-response behaviors.

Overall, these findings indicate that copper-based UFPs generated by spark discharge processes can trigger biologically relevant pulmonary effects at exposure levels comparable to polluted occupational environments, supporting their potential health relevance and the need for further mechanistic and in vivo correlation studies.

## Acknowledgements

The authors would like to thank all contributors and collaborators involved in the ULTRHAS project. We gratefully acknowledge the technical support provided by the Core Facility Genomics (Dr. Inti Alberto de la Rosa Velazquez) and the Core Facility Metabolomics and Proteomics (Dr. Agnese Petrera) at Helmholtz Munich.

## Author contributions

**Conceptualization:** Johannes Becker, Martin Sklorz, Sebastiano Di Bucchianico, Ralf Zimmermann. **Methodology**: Johannes Becker, Jana Pantzke, Svenja Offer, Anusmita Das, Martin Sklorz, Sebastiano Di Bucchianico. **Formal analysis and investigations**: Johannes Becker, Jana Pantzke, Svenja Offer, Anusmita Das, A. Mudan, Carsten Neukirchen, Martin Sklorz, Sebastiano Di Bucchianico. **Writing – original draft**: Johannes Becker. **Funding acquisition**: Sebastiano Di Bucchianico, Thomas Adam, Ralf Zimmermann. **Resources**: Thomas Adam, Ralf Zimmermann. **Supervision**: Martin Sklorz, Sebastiano Di Bucchianico, Ralf Zimmermann. **Project administration**: Thorsten Streibel, Sebastiano Di Bucchianico. **Visualization**: Johannes Becker, Sebastiano Di Bucchianico. **Writing – review & editing**: all authors.

## Funding

The research leading to these results received funding from the project ULTRHAS (ULtrafine particles from TRansportation – Health Assessment of Sources) under Grant Agreement No 955390 from the EU’s Research and Innovation program Horizon 2020. Views and opinions expressed are, however, those of the author(s) only and do not necessarily reflect those of the European Union or CINEA. Neither the European Union nor the granting authority CINEA can be held responsible for them. This research is funded by dtec.bw – Digitalization and Technology Research Centre of the Bundeswehr [project LUKAS and MORE]. Dtec.bw is funded by the European Union – NextGenerationEU.

## Data availability

Data are available upon reasonable request.

## Ethical approval

None

## Conflict of interest

The authors declare no conflict of interest.

## Notes

### Competing Interest Statement

The authors have declared no competing interest.

