## supplementary material for "Railway Catenary Sparking as a Source of Toxic Copper Ultrafine Particles: Evidence from Realistic In Vitro Inhalation Exposure"

Volcano Plots depicting the annotated genes of the RNA-seq analysis of the A549 cells and the 3D model exposed to SDG-UFP at 100 mL/min.

**Supplementary figure 1**

Differently expressed genes (DEGs) in the A549 model mapped by RNA-seq analysis (-$\log_{10}$ of p-value $\geq 1.3$; $\log_2$ FC $\geq \pm 0.58$).

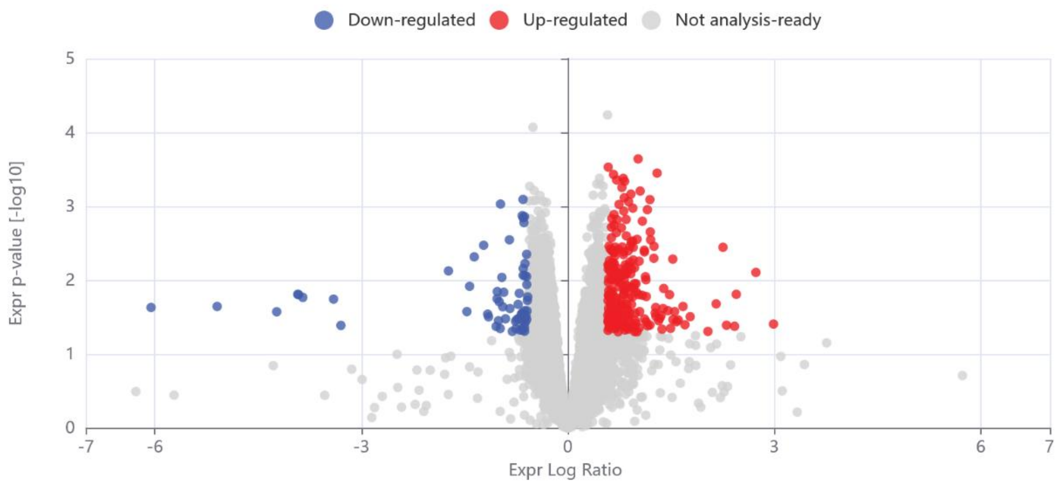


**Fig. S1** Volcano plot of the RNA-seq analysis in the A549 model highlighting the DEGs. Exposure of A549 yielded 240 DEGs upregulated (red) and 64 DEGs downregulated (blue) more than 1.5-fold over CA exposed cells. Data shown as $\log_2$ of FC expression (x-axis) and -$\log_{10}$ of p-value (y-axis) as mean of three independent experiments

**Supplementary figure 2**

Differently expressed genes in the Calu-3 and dTHP-1 cells from the 3D model mapped by RNA-seq analysis (-$\log_{10}$ of p-value $\geq$ 1.3; $\log_2$ FC $\geq \pm$ 0.58).

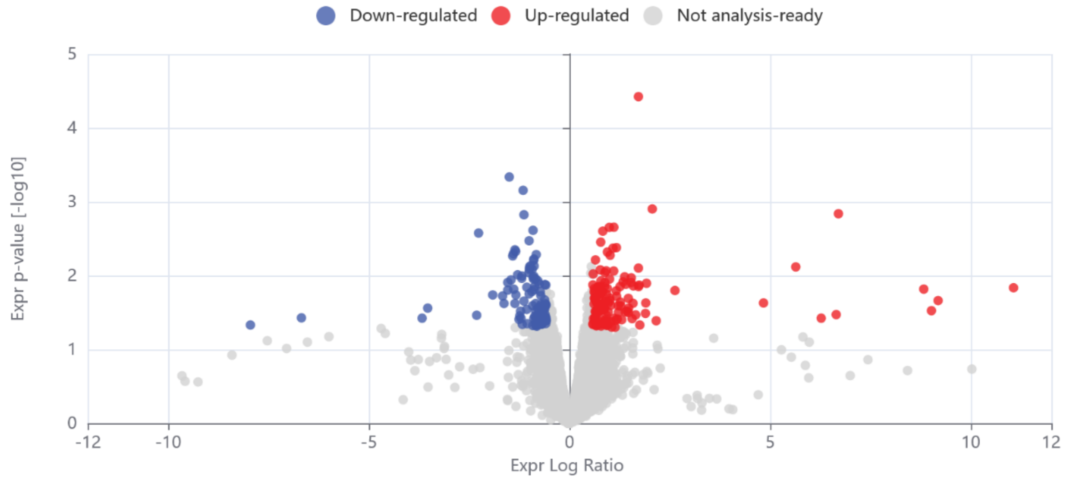


**Fig. S2** Volcano plot of the RNA-seq analysis in the apical 3D model highlighting the DEGs. Exposure of the apical Calu-3 and dTHP-1 cells yielded 55 DEGs upregulated (red) and 37 DEGs downregulated (blue) more than 1.5-fold over CA exposed cells. Data shown as $\log_2$ of FC expression (x-axis) and -$\log_{10}$ of p-value (y-axis) as mean of three independent experiments

**Supplementary figure 3**

Differently expressed genes in basolateral, endothelial EA.hy926 cells from the 3D model mapped by RNA-seq analysis (-$\log_{10}$ of p-value $\geq$ 1.3; $\log_2$ FC $\geq \pm$ 0.58).

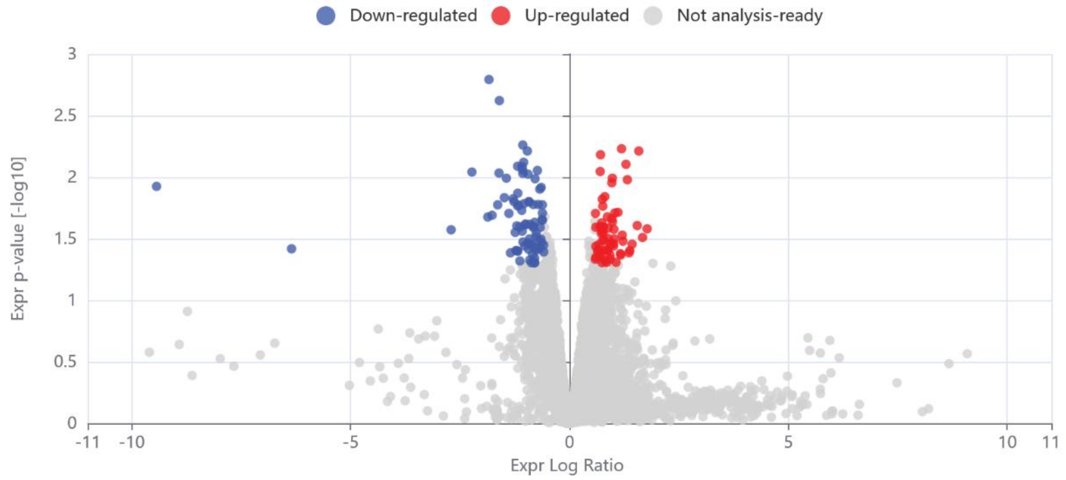


**Fig. S3** Volcano plot of the RNA-seq analysis in the basolateral 3D model highlighting the DEGs. Exposure of the basolateral EA.hy926 cells yielded 61 DEGs upregulated (red) and 77 DEGs downregulated (blue) more than 1.5-fold over CA exposed cells. Data shown as $\log_2$ of FC expression (x-axis) and -$\log_{10}$ of p-value (y-axis) as mean of three independent experiments
